# Nicotine dependence (trait) and acute nicotinic stimulation (state) modulate attention but not cognitive control: converging fMRI evidence from Go-Nogo and Flanker tasks

**DOI:** 10.1101/770651

**Authors:** E. Lesage, M.T. Sutherland, T.J. Ross, B.J. Salmeron, E.A. Stein

## Abstract

Cognitive deficits during nicotine withdrawal may contribute to smoking relapse. However, interacting effects of chronic nicotine dependence and acute nicotine withdrawal on cognitive control are poorly understood. Here, we examine the effects of nicotine dependence (trait; smokers versus non-smoking controls), and acute nicotinic stimulation (state; administration of nicotine and varenicline, two FDA-approved smoking cessation aids, during abstinence), on two well-established tests of cognitive control, the Go-Nogo task and the Flanker task, during fMRI scanning. We compared performance and neural responses between these four pharmacological manipulations in a double-blind, placebo-controlled crossover design. As expected, performance in both tasks was modulated by nicotine dependence, abstinence and pharmacological manipulation. However, effects were driven entirely by conditions that required *less* cognitive control. When demand for cognitive control was high, abstinent smokers showed no deficits. By contrast, acutely abstinent smokers showed performance deficits in easier conditions and missed more trials. Go-Nogo fMRI results showed decreased inhibition-related neural activity in right anterior insula and right putamen in smokers and decreased dorsal anterior cingulate cortex activity on nicotine across groups. No effects were found on inhibition-related activity during the Flanker task, or on error-related activity in either task. Given robust nicotinic effects on physiology and behavioral deficits in attention, we are confident that pharmacological manipulations were effective. Thus, findings fit a recent proposal that abstinent smokers show decreased ability to divert cognitive resources at low or intermediate cognitive demand, while performance at high cognitive demand remains relatively unaffected, suggesting a primary attentional deficit during acute abstinence.

## INTRODUCTION

Tobacco addiction remains the leading cause of preventable death in the world^1^. The vast majority of smokers express a strong desire to quit, but most quit attempts fail within a week^2^. An important cause of relapse is the nicotine withdrawal syndrome (NWS), which is characterized by affective^3,4^, reward^5–7^ and cognitive deficits^8,9^. Cognitive control deficits have been hypothesized to be a key symptom of drug withdrawal^10,11^, both because they impact normal daily functioning and because they induce a deficit state making it more difficult to remain abstinent.

Nicotine is a nonselective nicotinic acetylcholine receptor (nAchR) agonist, acting on multiple neurotransmitter systems, including dopamine (DA), norepinephrine (NE), glutamate and GABA^12^. Nicotine is an indirect DA agonist, binding to α4β2 nArchRs on midbrain DA neurons and stimulating the mesocorticolimbic (MCL) system, which consists of striatal, limbic and prefrontal terminals of these midbrain DA neurons^13^. Nicotine’s actions on different neurotransmitter systems can interact with each other. For example, the NE system targets overlapping prefrontal circuitry to the MCL system and modulates attention^14^, and nicotine’s glutaminergic effects modulate descending corticostriatal pathways^15^. Nicotine’s reinforcing properties derive from its action upon the dopaminergic MCL system. Dependence is thought to develop through chronic stimulation of this system, leading to neuroplastic changes in MCL circuitry that downregulate DA levels^11,16^. This hypodopaminergic withdrawal state that characterizes drug dependence is one prominent mechanistic hypothesis through which abstinence from nicotine could impair smokers’ cognitive control.

In line with this mechanistic hypothesis, effective pharmacological smoking cessation aids such as nicotine replacement therapy (NRT) and varenicline (Chantex®), predominantly target α4β2 nAchRs. The latter serves as a partial agonist at these receptors, acting as a weak agonist in the absence of nicotine, and as a partial antagonist in the presence of nicotine^17^. While the in vitro receptor binding mechanisms of these drugs and their efficacy at a clinical level is established^18–20^, it is less clear how their systems-level neurobiological mechanisms affect the cognitive and affective deficits seen in the human NWS. Moreover, these drugs are administered clinically in both the nicotine dependent (addiction trait) condition as well as in the acute withdrawal and prolonged abstinent state. How their neurobiological mechanisms of action are biased under these disease cycles and how that might affect their clinical efficacy are poorly understood. To this end, we investigate the effects of acute nicotinic receptor stimulation through a nicotine patch and varenicline pill interaction on cognitive control mechanisms in both smokers and non-smoking control participants.

We use two tasks to probe cognitive control and error monitoring: the Go-Nogo task and the Eriksen Flanker task^21^. These have previously characterized cognitive control deficits in substance abuse, including cocaine, alcohol and nicotine^22,23^. Both tasks tax inhibitory control over a dominant response tendency and largely recruit the same neural networks^24–26^. The Go-Nogo task requires the inhibition of a prepotent motor response (especially when Nogo trials are rare, as in this study), while in the Flanker task response dominance elicited by an irrelevant task dimension must be suppressed^26^.

Neuroimaging studies in substance dependent populations have consistently reported hypoactivation in frontal cortical areas associated with cognitive control^22^, notably the dorsal anterior cingulate cortex (dACC), dorsolateral prefrontal cortex (DLPFC), and right inferior frontal gyrus (IFG). Administration of a DA antagonist decreases neural activity in these regions and cognitive control performance in both smokers and non-smokers^27^. This hypofrontality is consistent with the DA deficiency model of addiction^28^ and generalizes to other substance use disorders, including cocaine addiction^29^.

Behavioral evidence for cognitive control deficits in nicotine-dependent populations is mixed, with some studies reporting deficits^30,31^, while others do not^32,33^. The inconsistency in the literature may be partially attributable to the drug state of the cohorts used; most studies compared nicotine-sated smokers with non-smokers while others required smokers to be abstinent^23,30,31^. To the best of our knowledge, no studies have directly examined the effects of nicotine dependence on cognitive control behavior and neural mechanisms in both the nicotine withdrawal (state) and nicotine sated conditions (trait), even though these comparisons are crucial to understanding the NWS and its role in treatment success or more often, relapse.

The current study overcomes these limitations by manipulating both the current state (12 hours abstinent vs. acutely sated) and trait (smokers vs. non-smokers). We examined the effects of nicotine patch and varenicline pill (both alone and in combination to mimic how these drugs are used clinically) using a Go-Nogo and a Flanker task as probes of cognitive control. Both smokers and non-smokers performed these tasks during fMRI acquisition; the 12-hour abstinent manipulation in smokers was intended to mimic the first day of a quit attempt, which is when smokers are most vulnerable to relapse ^34^. We hypothesize that abstinent smokers will perform worse than non-smokers on these tasks, specifically in task conditions that require higher cognitive control, and that this deficit will be alleviated, at least in part, by nicotine and, to a lesser extent, varenicline. In addition, we expect that neural responses during successful and failed inhibition will be affected by chronic nicotine dependence trait and acute nicotinic stimulation in a manner reflecting their effects on performance.

## METHODS

### Participants

Participants were 24 smokers (12 female) and 20 non-smokers (10 female). Written informed consent was obtained from the NIDA-IRP Institutional Review Board. Participants were right-handed, between the ages of 18-55, and had no reported history of neurological or psychiatric disorders or current or past substance dependence (other than nicotine in smokers). Non-smokers reported no history of daily nicotine use and no smoking within the last 2 years. Groups were matched on gender and ethnicity (Table S1). Smokers were older than controls (t_41.32_=2.12, p<0.05), and age was therefore included as a covariate in all between-group analyses. Data from one male non-smoker were excluded due to consistently poor behavioral performance and excessive head motion.

### Study design

Participants underwent 6 scanning sessions as part of a fully counterbalanced, two-drug, double-blind, placebo-controlled study. At three points during a varenicline administration regime, (i.e. pre-pill, 2 weeks varenicline pill, 2 weeks placebo pill), participants were scanned twice, once wearing a nicotine patch and once wearing a placebo patch. Results from the four completely counterbalanced sessions are reported herein. The reported data are part of a larger study that also probed effects of nicotine and varenicline on reward anticipation^5^, economic decision making^35^, and emotional reactivity^4,36^. For detailed information on the experimental design and MRI acquisition parameters see Supplementary Methods.

### Go-Nogo and Flanker tasks

Participants carried out four 5-minute blocks (300 trials each) of a Go-Nogo task with alternating “X” and “Y” stimuli^25^ (see Supplementary Methods and Figure 1A). Trials were spaced 1s apart, with the relatively infrequent Nogo trials (100 out of 1200 trials; 8.3%) temporally jittered to optimally estimate the BOLD response. Participants also performed four 9-minute 130-trial runs of a speeded version of the Flanker task (see Supplementary Methods and Figure 2A) with an individualized response deadline to ensure an adequate difficulty level and error rate^36^.

**Figure 1.**
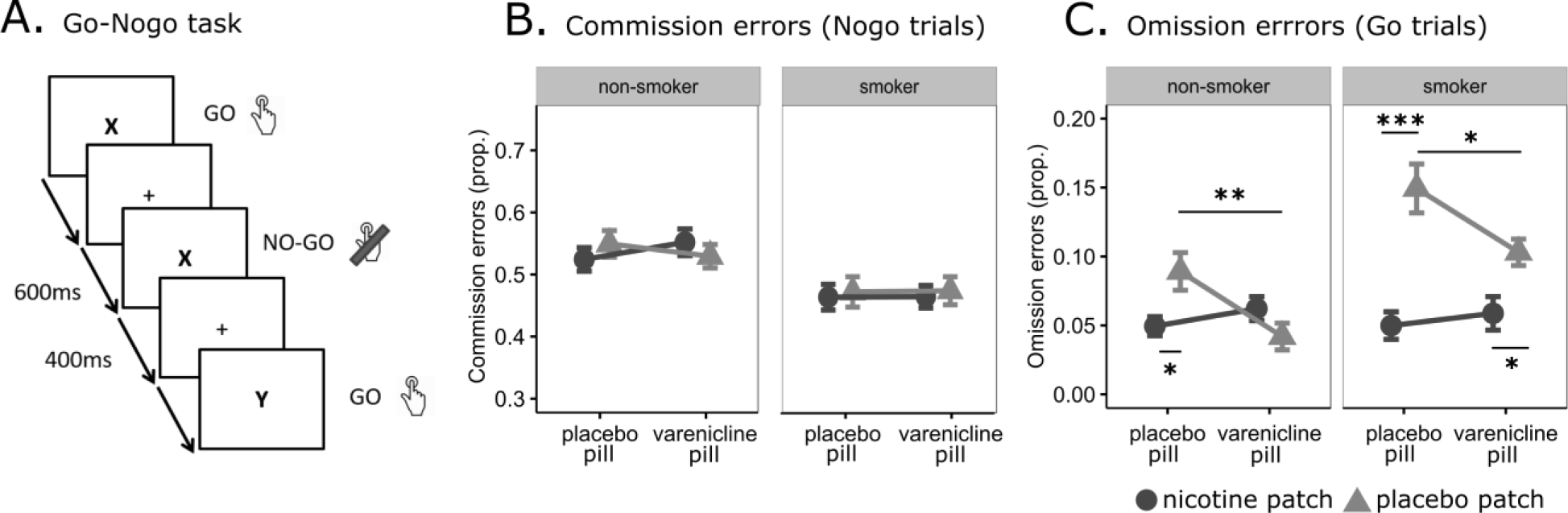
Go Nogo task. A. Trial timing of the task. Nogo trials are modeled against the implicit baseline which consists of Go trials and fixation between trials. The timing of Nogo trials is temporally jittered so that these trials can be modeled. B-C: Go-Nogo performance. Error rate on Nogo (B) and Go (C) trials. Error bars indicate +/− 1 standard error of the mean. * p<0.05, ** p<0.01, *** p<0.001.

**Figure 2:**
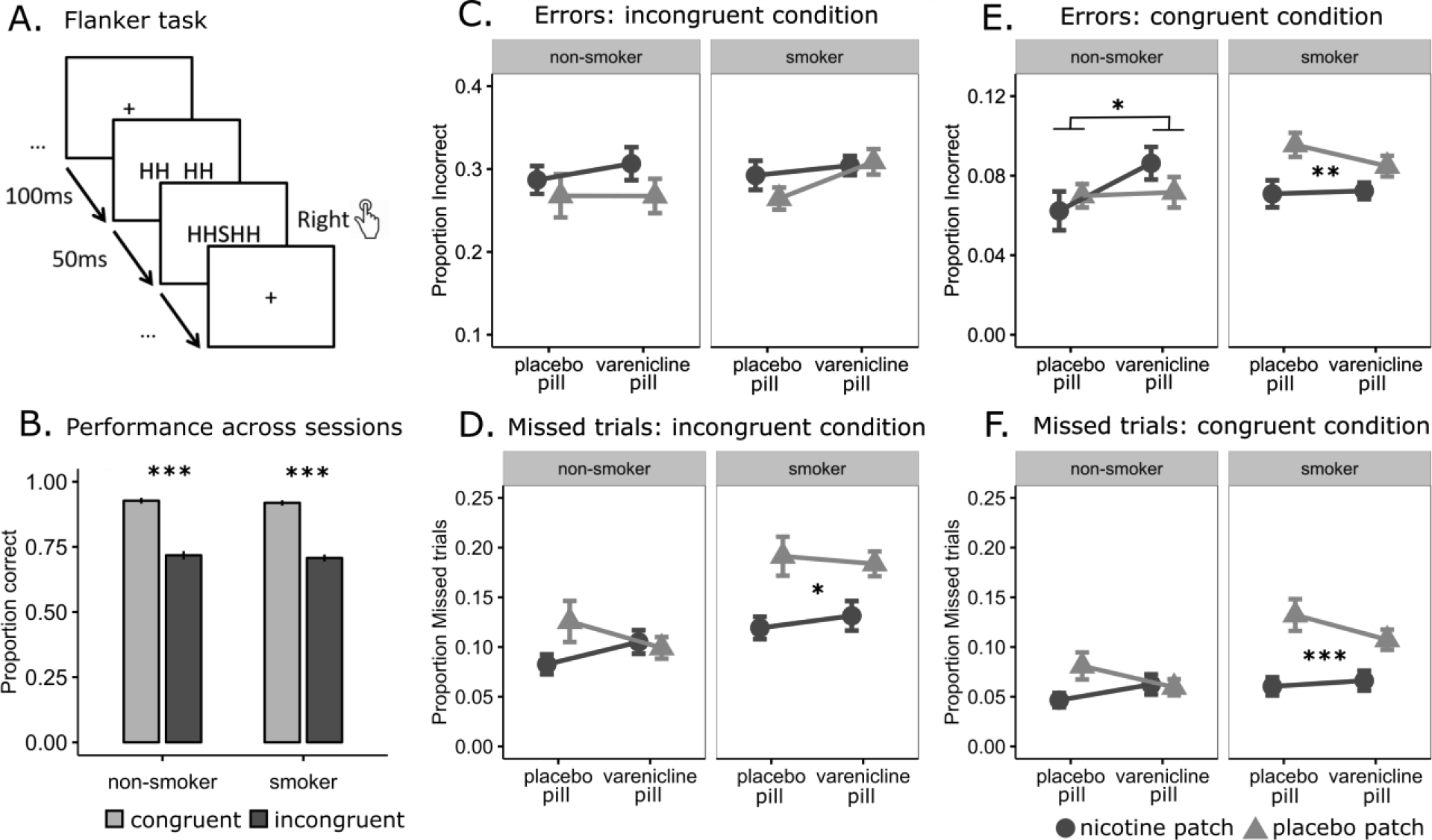
Flanker task A. Trial timing of the task. Ellipses indicate variable temporal jitter, enabling statistical modeling of each trial against the implicit baseline consisting of fixation periods between trials. B-F: Behavioral performance. B. Performance across sessions. C-D. Performance on Incongruent trials. E-F. Performance on congruent trials. Error bars indicate +/− 1 standard error of the mean. * p<0.05, ** p<0.01, *** p<0.001.

### Behavioral analyses

Analyses were performed in R (www.r-project.org/) using packages afex and phia. For the Go-Nogo task, we performed mixed-model ANOVAs on two dependent variables: the error rate on Nogo trials (commission errors, indicating a failure of response inhibition), and the error rate on Go trials (omission errors, indicating attention lapses). For the Flanker task, we performed mixed-model ANOVAs on four dependent variables: the error rate on congruent and on incongruent trials (indicating a failure to overcome a prepotent response), and the missed trial rate on both trial types (the latter three indicating a lack of attention). We compared performance on incongruent and congruent trials across sessions as a manipulation check. Independent variables were the between-subject variable GROUP (smokers vs. non-smokers), and the within-subject variables NICOTINE (nicotine patch vs. placebo patch) and VARENICLINE (varenicline pill vs. placebo pill), and age as a covariate. Interactions were followed up with within-group ANOVAs using NICOTINE and VARENICLINE as independent variables.

### Imaging analyses

#### Preprocessing

Imaging analyses were carried out in AFNI^37^. Functional scans were preprocessed using standard methods: slice-time correction, image alignment to MPRAGE, motion correction, spatial registration to the Talairach template and smoothed to 8mm FWHM.

#### Single subject analysis

Three Go-Nogo regressors of interest were modeled against the implicit baseline: correctly executed Nogo trials, incorrectly executed Nogo trials, and incorrectly executed Go Trials. Correctly executed Go trials were included into the implicit baseline. Two contrasts of interest were calculated: INHIBITION [Nogo Correct (–) implicit baseline] and ERROR [Nogo Incorrect (–) Nogo Correct]. Four Flanker regressors of interest were modeled against the implicit baseline: correct and incorrect responses for congruent and incongruent trials. Two contrasts of interest were estimated: INHIBITION [Incongruent Correct (–) Congruent Correct] and ERROR [Incongruent Incorrect (–) Incongruent Correct]. We excluded (censored) timepoints where Euclidian displacement between successive frames exceeded 0.3 mm or where the DVARS^38^ exceeded 1.3. For the Flanker task, missed responses were modelled separately as a regressor of no interest. For the Go-Nogo task, timepoints where responses were absent for 10 trials or more were similarly modeled separately. First level design matrices for both tasks included the regressors of interest and their temporal derivative, polynomials for each block, 6 regressors that modeled head motion and regressors of no interest capturing the censored timepoints.

#### Group level Analysis

*Average activity patterns* for INHIBITION and ERROR contrasts were computed with t-tests over subjects’ beta weight maps (averaged over sessions). Results were whole-brain FWE corrected (alpha<0.05, voxel-wise p<0.001, cluster-size 22 voxels).

*Group and drug effects* for both contrasts were examined with mixed ANOVAs (between-subject factor GROUP, within-subject factors NICOTINE and VARENICLINE, age as covariate). Significant GROUP interactions were followed by within-group analyses using NICOTINE and VARENICLINE as factors. We constructed a functional small-volume mask of interest, derived separately for each task from an OR mask of the INHIBITION and the ERROR contrasts (see Supplementary Methods). Results within this volume of interest were corrected at FWE alpha<0.05, as determined by a 3dClustSim algorithm in AFNI^37^ which indicated a voxelwise threshold of p<0.01 with a minimum cluster size of 24 for the Go-Nogo task and 22 for the Flanker task.

## RESULTS

### Behavioral results

#### Go-Nogo performance

The rate of commission errors, the failure to suppress a prepotent response, was unaffected by GROUP, NICOTINE, VARENICLINE or their interactions (Figure 1B). By contrast, omission errors, missed responses on Go trials, showed significant NICOTINE (F(1,123)=33.72, p<0.001), NICOTINE*GROUP (F(1,123)=6.22, p=0.014) and NICOTINE*VARENICLINE (F(1,123)=11.55, p<0.001) effects. Specifically, smokers in the acute abstinence (placebo patch) condition committed more omission errors than under the nicotine patch condition, and the nicotine effect interacted with varenicline in accordance to their known drug-drug pharmacological interactions^17^. Although nicotine slightly reduced omission rate in nonsmokers, its effect on omission rate was larger in smokers than non-smokers (Figure 1C).

#### Flanker performance

Both smokers (F(1,161)=77.18, p<0.001) and non-smokers (F(1,126)=56.11, p<0.001) performed better in the congruent than the incongruent condition, with no group difference across sessions (CONDITION*GROUP, F(1,287)=0.006, p=0.939, Figure 2B). Examining accuracy per condition, no main or interaction effects of GROUP, NICOTINE, or VARENICLINE were found in the incongruent condition (Figure 2C). By contrast, in the congruent condition, a main effect of NICOTINE (F(1,123)=5.68, p=0.018), and trends for VARENICLINE (F(1,123)=3.55, p=0.062), GROUP*VARENICLINE (F(1,123)=2.77, p=0.099), and NICOTINE*VARENICLE (F(1,123)=3.24, p=0.074) were identified (Figure 2E). These patterns were driven by worse performance in smokers without nicotine (F(1,69)=9.49, p=0.003) and better performance with varenicline in non-smoker group (F(1,54)=4.46, p=0.039). Both groups missed more trials in the incongruent than the congruent conditions (F(1,287)=11.45, p<0.001), missed fewer trials with the nicotine patch (F(1,287)=31.20, p<0.001), with this nicotinic effect smaller in the presence of varenicline pill (NICOTINE*VARENICLINE: F(1,287)=5.95, p=0.015) and tended to be larger in smokers (NICOTINE*GROUP: F(1,287)=2.80, p=0.099; Figures 2D and 2F). These patterns were explained by a strong NICOTINE main effect in smokers (F(1,69)=19.10, p<0.001), and a weaker NICOTINE effect in non-smokers (F(1,54)=5.84, p=0.019).

In sum, for both cognitive control tasks, we failed to find effects of smoker (GROUP) or acute nicotinic stimulation (NICOTINE or VARENICLINE) in conditions that require greatest cognitive control (Nogo condition and Incongruent condition). In contrast, we do find robust DRUG and smoker (GROUP) effects in the *less* demanding control conditions and in the rate of missed responses. This suggests that acute nicotine withdrawal impairs low level, attentional processes rather than higher level cognitive control, and that this impairment is ameliorated by nicotinic stimulation.

### Neuroimaging results

#### Go-Nogo task

The INHIBITION task map (Figure 3A) shows robust activity in the salience network^39^, encompassing bilateral inferior frontal gyrus, anterior insula, and dorsal anterior cingulate cortex (dACC), stretching out into presupplementary motor areas (pre-SMA), as well as the striatum and thalamus. Smaller areas in medial frontal cortex and precuneus showed less activation during Nogo trials than during Go trials. These patterns are in line with meta-analysis of Go-Nogo tasks ^25^. In the ERROR contrast (Figure 3B), regions of the dACC and pre-SMA, left dorsal striatum, middle frontal gyrus and cuneus showed increased BOLD activation during commission errors compared to correctly withholding a response.

**Figure 3.**
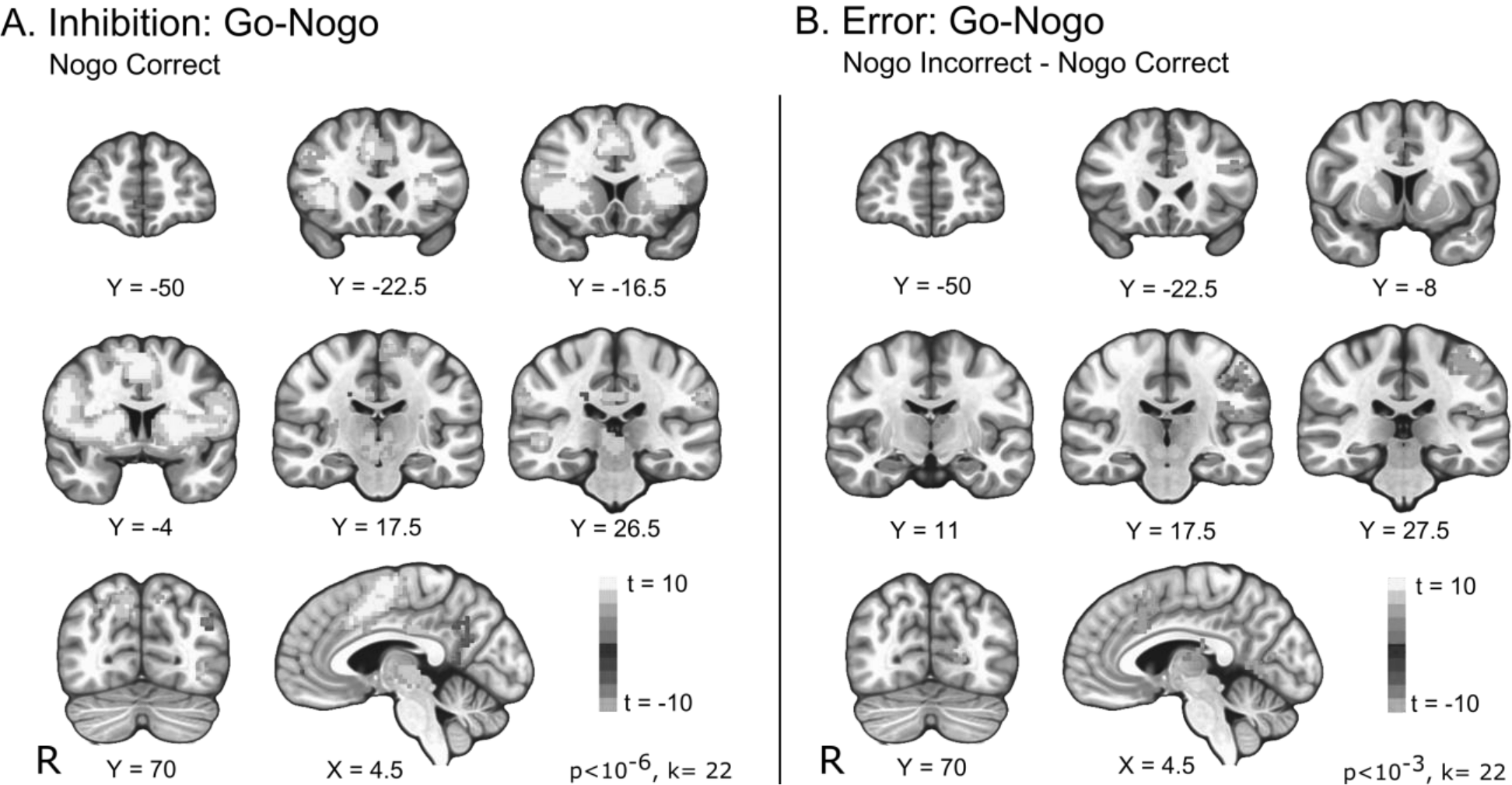
Task maps for the Go-Nogo contrasts. A. Brain areas with greater (orange) or smaller (blue) response to successfully inhibiting a prepotent response compared to baseline (correctly executed Go trials). Image whole-brain corrected at p<10^−6^ voxelwise for display purposes, see Figure S3 for image corrected at p<10^−3^. B. Brain areas that showed greater (orange) response to failed inhibitions than to successful inhibitions on Nogo trials. Whole-brain corrected p<10^−3^ voxelwise, cluster size 22, FWE<0.05. Radiological orientation: right is presented on the left.

Effects of GROUP, NICOTINE and VARENICLINE were assessed with mixed ANOVA for both contrasts, FWE corrected within the task mask of interest (Figures 4A and 4B). There was a GROUP effect in the right anterior insula and right putamen: smokers showed less activity than non-smoking controls (Figure 4A). In addition, there was a main effect of NICOTINE within a region on the border of dACC and pre-SMA that was less active under the nicotine patch compared to the placebo patch condition (Figure 4B). No session effects were found for the ERROR contrast.

**Figure 4.**
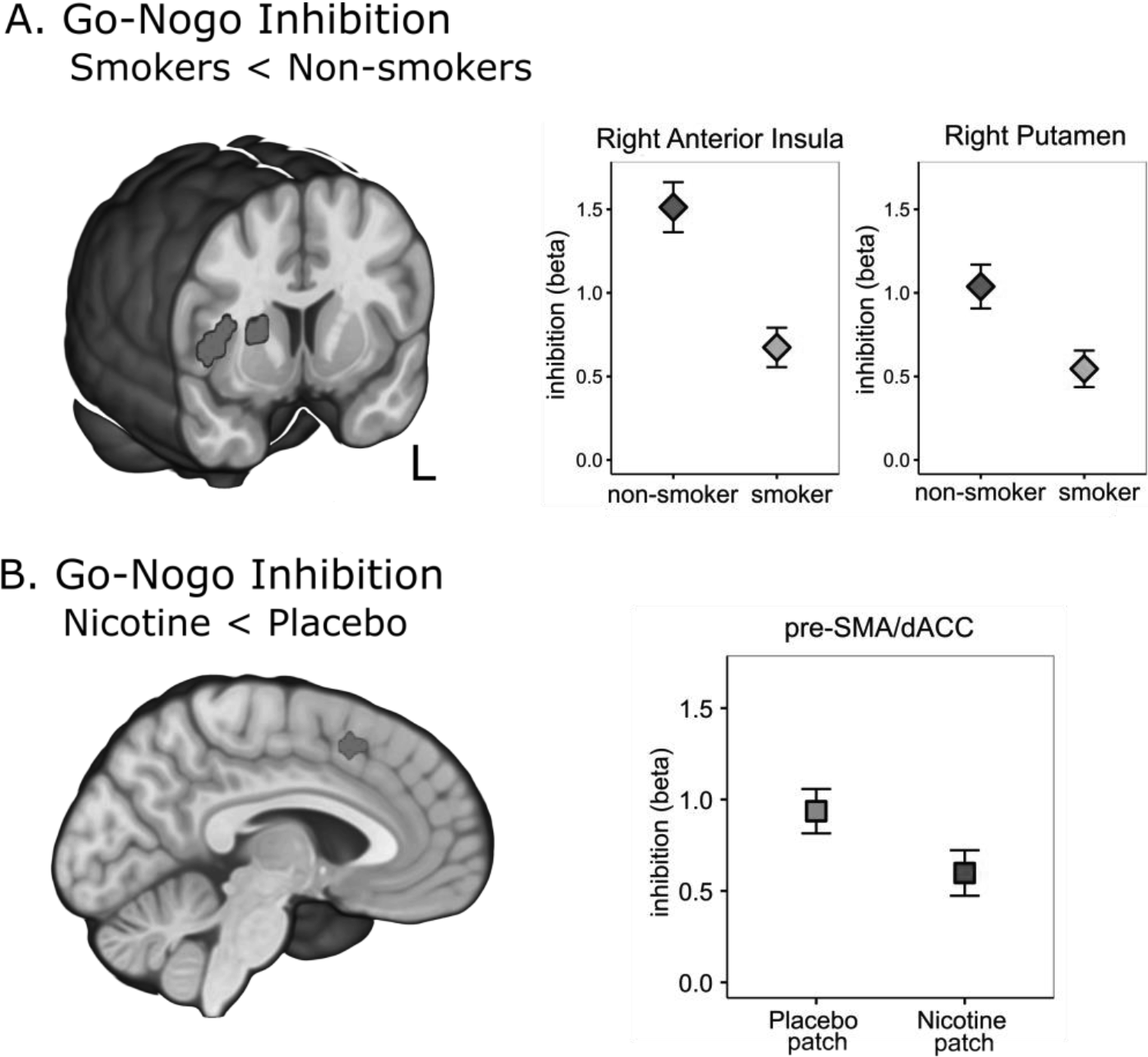
A. Smokers show hypoactivation of right anterior insula and right putamen during correct inhibition of a prepotent response compared to non-smokers. B. Across groups, nicotine lowered inhibition-related activity in the superior frontal gyrus (pre-SMA/dACC). Both panels: Activations FWE corrected within the mask of interest (p<0.01 voxelwise, minimum cluster size 24 voxels in Go-Nogo task, 22 voxels in Flanker task). Extracted regression weights and error bars (SEM) presented to aid interpretation only; no statistical inference should be drawn. Radiological orientation: right is presented on the left.

#### Flanker task

The INHIBITION contrast (across all sessions) revealed increased activation during correct incongruent trials compared to correct congruent trials in bilateral AI and ACC/pre-SMA (nodes of the salience network^39^, as well as bilateral dlPFC and left superior parietal cortex (Figure 5A), consistent with a previous meta-analysis of Flanker tasks^40^. In contrast, the right inferior parietal, left occipital and left thalamus showed greater response to congruent compared to incongruent conditions. The ERROR contrast identified a robustly activated network including bilateral AI, dACC/pre-SMA and SMA, bilateral thalamus, DLPFC, superior parietal lobule, and occipital cortex (Figure 5B). The mixed ANOVA examining effects of GROUP, NICOTINE and VARENICLINE within these masks did not yield any significant session effects for either task contrast.

**Figure 5.**
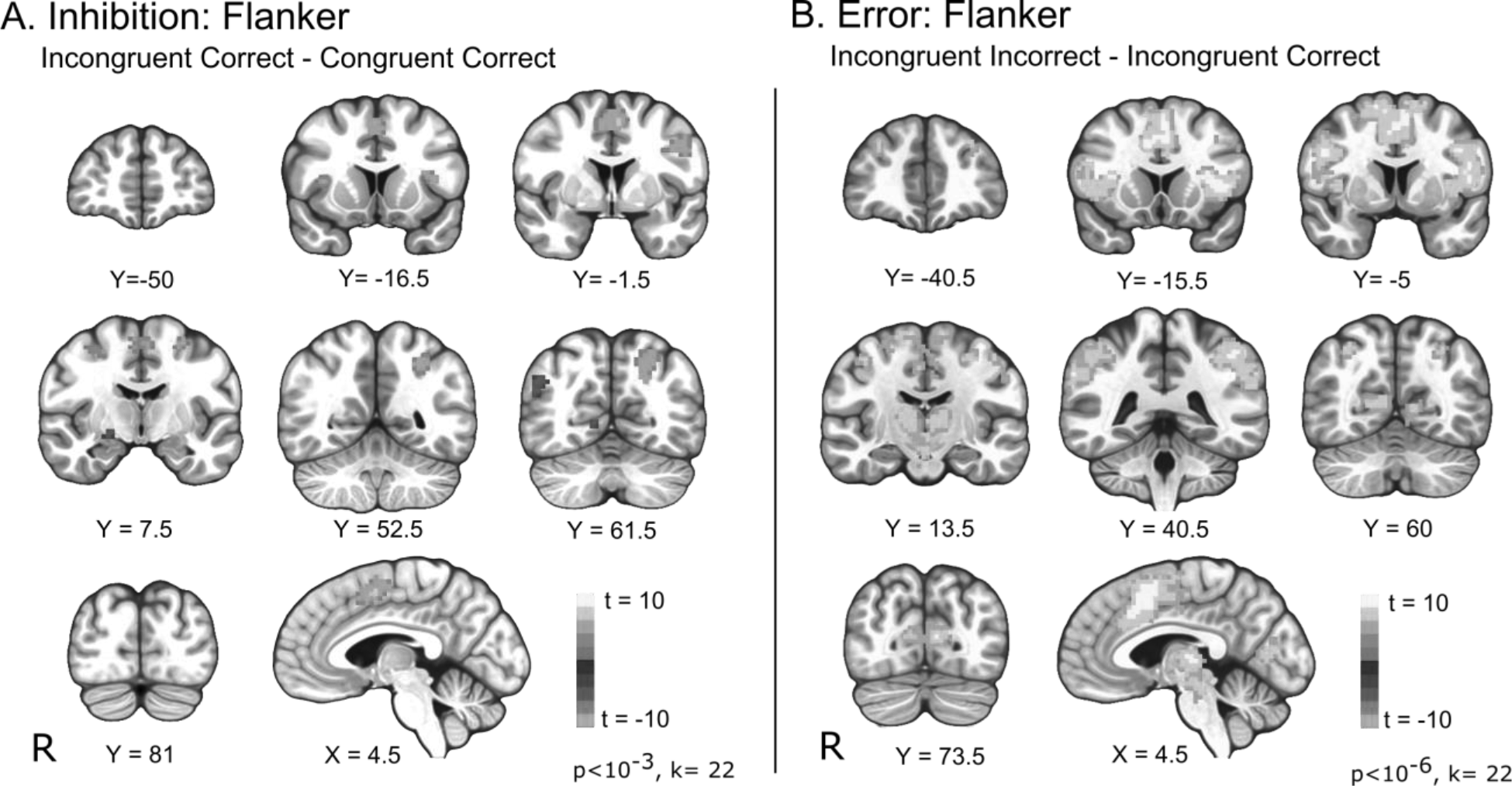
Task maps for Flanker task. A. Brain areas that showed more (orange) or less (blue) activity in response to correctly executed incongruent trials than to correctly executed congruent trials. Whole-brain corrected at FWE <0.05 (p<0.001 voxelwise, cluster size 22). B. Brain areas that showed greater activity during the incorrect execution of an incongruent trial than to the correct execution of an incongruent trial. Figure whole-brain corrected p<10^−6^ voxelwise for display purposes; see Figure S4 for figure corrected at p<10^−3^. Radiological orientation: right is presented on the left.

## DISCUSSION

The nicotine withdrawal syndrome (NWS), characterized by dysregulated affective and cognitive processing, has been hypothesized to be the principal factor in smoking cessation failures. As such, a better understanding of the neurobiological mechanisms underlying the NWS and how pharmacotherapies might help alleviate specific withdrawal signs linked to relapse is critically needed. In this study, we hypothesized that acute nicotine withdrawal in smokers would disrupt cognitive control processing, which would be reversed in the presence of two commonly used pharmacotherapeutic agents: nicotine patch and/or varenicline pill. We employed two cognitive control tasks to test this hypothesis: the Go-Nogo and the Flanker task. Contrary to our hypothesis, we did not find effects on performance in demanding conditions requiring inhibitory control of either task during abstinence. However, we did find the hypothesized drug and condition (abstinence vs. satiety) effects on measures of attention and on the number of missed trials (i.e. omissions) in both tasks. That is, our behavioral results indicate deficits in sustained attention in abstinent, but not sated smokers, that are alleviated with nicotinic receptor stimulation. Imaging analyses during the successful withholding of an inappropriate response in the Go-Nogo showed lower right anterior insula and putamen activity in smokers independent of state, and decreased dACC/pre-SMA activity in the presence of a nicotine patch.

Our results show that nicotine dependence trait and nicotinic stimulation effects (state) were present in conditions requiring the *least* cognitive control. Notably, patterns were strikingly similar across the two tasks, adding confidence to this somewhat counter-intuitive finding. In the Go-Nogo task, abstinent smokers failed to execute responses when they should have, i.e. errors of omission. In the Flanker task, 12hr nicotine deprived smokers were more likely to fail to press the correct button when flankers primed the correct location. Moreover, missed responses were more prominent in abstinent smokers for both congruent and incongruent conditions, with this deficit marginally stronger in the congruent (less demanding) condition. In all cases, these deficits were alleviated with the administration of nicotine, following which performance was similar to that of non-smokers. Varenicline, a partial agonist at the α4β2 nicotinic receptors, significantly improved performance only in the Go-Nogo task. Generalizing across tasks, abstinent smokers appear to fail to allocate appropriate attentional resources at low levels of cognitive demand. Rather than a blanket inability to engage cognitive and attentional resources, the deficit appears specific to conditions where only relatively minimal attention is required. The robust effects on attention that we report are in line with literature on attention-enhancing effects of nicotine^14,41^ and problems with sustained attention in smokers^42–45^.

In contrast, we did not find behavioral effects in task conditions requiring high cognitive control. While it is possible that such effects exist but were undetectable in this sample, this seems unlikely given the presence of clear drug effects at the physiological level and in the control conditions, the reproducibility using two independent, well characterized tasks, and effects on other tasks performed in the same cohort and in the same session, including reward processing, emotional processing and reversal learning^4,5,35,46^. We are therefore confident that the imposed pharmacological manipulations exerted appropriate pharmacodynamic effects. Moreover, the absence of cognitive control deficits is not exceptional in the literature. Evidence of nicotine dependence on cognitive control has been mixed. Some studies report worse cognitive control performance in smokers^30,31,47^, while others did not^33,48,49^. Studies manipulating satiety within-subject similarly reported mixed results for cognitive control tasks^42,44^.

One way to interpret our results is to consider the varying demand hypothesis proposed by Fedota et al^50^. Using a Flanker task with three levels of difficulty, the authors found that smokers and non-smokers’ performance and brain activity only differed at intermediate levels of congruency/difficulty. At the highest and the lowest levels of inhibitory demand, both groups performed equally well and similarly recruited task-relevant brain areas. Thus, rather than a generalized reduction in cognitive control capacity, it may be that abstinent smokers’ deficits lie in the inability to allocate cognitive and attentional resources in response to slightly increased demand. That is, when inhibitory demand is clearly high, abstinent smokers can recruit the necessary resources, but when inhibitory demand is perceived to be low or intermediate, these resources are not adequately recruited and performance drops. Smokers, even when abstinent, might therefore be just as capable of engaging cognitive control resources as non-smokers, but simply fail to do so at similar levels of environmental challenge. This interpretation may also help shed some light on inconsistencies in the literature. Subtle difference in the design of the tasks, or the recruited population (especially individual differences in cognitive capacity), may cause inhibitory control (Nogo, Incongruent) conditions to be perceived either as intermediately demanding (where one would expect group differences) or as very demanding (where these differences are not expected).

Imaging results showed lower right anterior insula and putamen activity in smokers during the successful withholding of an inappropriate response and decreased dACC/pre-SMA activity in the presence of a nicotine patch. Reduced engagement of dependent smokers’ frontal and striatal task-relevant brain areas is in line with neuroimaging studies on nicotine dependence^22,31^ and other substance use disorders^29^. Interestingly, the right anterior insula has been specifically implicated in allocating cognitive resources by monitoring performance and environmental demands^51,52^. Across groups, the dACC/pre-SMA, which is central to processing salience, attention, and inhibitory control in both tasks^25,39,40^, showed less brain activity in the nicotine patch compared to the placebo patch condition, indicating that less recruitment of this area is needed to achieve the same performance. Previous neuroimaging studies have similarly reported decreased dACC activity during a cognitive control task after smoking^53^ and in superior and middle frontal gyrus during an attentional task on nicotine patch^41^. Both studies were performed in smokers and results suggested that nicotinic effects were related to alleviation of withdrawal. In the current study, however, we were able to compare smokers and non-smokers, and did not find a group difference. The dACC/pre-SMA downregulation by nicotine may therefore reflect increased processing efficiency rather than an alleviation of decreased efficiency.

Our results have potential clinical implications. Manipulating the acute abstinent state in the smoker group allowed us to look at how cognitive control, as manifest using two response inhibition tasks, Go-Nogo and Flanker, and their neural signatures, are impacted by nicotine and varenicline. This is crucial to assess these drugs’ ability to alleviate cognitive deficits reported during the NWS. Rather, we show that smokers’ ability to pay attention to relatively easy tasks is diminished when abstaining from nicotine. Speculatively, newly abstinent smokers may be able to suppress the urge to smoke at the time and in the context they anticipate will be difficult but may be more vulnerable for relapse in contexts of lower perceived risk. It is encouraging that the deficits found were largely alleviated with the administration of a nicotine patch or (in the case of the Go-Nogo task) varenicline pill. This highlights the role smoking-cessation medication can play in supporting a quit attempt. Our results also support the use of cognitive behavioral approaches to enhance smokers’ awareness of the need to remain vigilant in seemingly less challenging situations.

This study is not without limitations. First, our fMRI design was optimized to test for differences in cognitive control, not sustained attention. In line with our behavioral results, few differences associated with successful (inhibition contrasts) and failed (error contrasts) trials were found. Our imaging tasks, particularly the Go-Nogo task, were not designed to test for differences in the less demanding control conditions, where behavioral effects did occur. We can speculate that group and session differences might have been found there, as they were behaviorally. Second, there is a chance that the modest sample size may have precluded observing a group or pharmacological effect. However, it is worth noting that our sample size is comparable or larger than those in previous studies^5,30,31,47^, and that the current sample showed robust behavioral and imaging effects on other tasks such as probabilistic reversal learning^35^, the monetary incentive delay task^5^ and an emotional reactivity task^4,36^, as well as on our measures of attention. Therefore, even if specific effects on inhibition were missed, these would likely have been very subtle.

Future investigations should further investigate smokers’ inability to allocate cognitive and/or attentional resources at lower or intermediate levels of difficulty. Moreover, effects on sustained attention, putatively mediated through nicotinic effects on norepinephrine signaling, should be factored into these designs, so that the contributions of these distinct effects might be teased apart.

## Notes

**Funding and disclosure:** This work was sponsored by the National Institute on Drug Abuse, Intramural Research Program, National Institutes of Health, US Department of Health and Human Services. EL is supported by the Flemish Fund for Scientific Research (FWO) grant FWO16/PG3/032. MTS was in part supported by National Institute on Drug Abuse grant K01DA037819, National Institute on Drug Abuse grant R01DA041353, and National Institute on Minority Health and Health Disparities grant U54MD01239 (sub-project 5378). The authors declare that there are no conflicts of interest to disclose.

