## Supplemental Materials for "Nicotine dependence (trait) and acute nicotinic stimulation (state) modulate attention but not cognitive control: converging fMRI evidence from Go-Nogo and Flanker tasks"

### METHODS

#### Study design

This study was a randomized, double-blind, placebo-controlled, crossover design involving two drugs: varenicline pills and nicotine patches. Subjects completed a total of 6 fMRI sessions: 2 baseline sessions (pre-pill), and 2 sessions each after a varenicline or placebo pill regimen. In each of the 3 varenicline conditions, the two sessions were randomized to either nicotine or placebo patch. Randomization was maintained by the study physician, while researchers, technicians, and participants remained blinded. This is in accordance with the protocol described in previous publications[1–4].

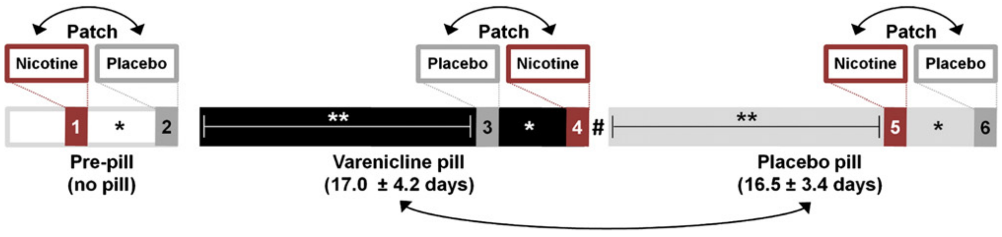


<u>Figure S1.</u> **Study design**. The current study reports data from the four completely counterbalanced sessions (sessions 3-6). Varenicline and placebo pill sessions were separated by more than two weeks (**). Nicotine and placebo patch scans were separated by an average of 2.9 ± 1.7 days (*). No washout interval separated medication periods (#). Double-headed arrows explain indicate randomized and counterbalanced order of the sessions. Figure reproduced with permission from Sutherland et al. [3].

#### Drug regimen

Varenicline and placebo pills were administered in accordance with standard guidelines (http://www.pfizer.com/products) over a ~17-day schedule. The varenicline regimen began at an 0.5 mg once-daily dosage for days 1-3, stepped up to 0.5 mg twice daily at days 4-7, and remained at 1 mg twice daily for days 8-17. Active and placebo medication appeared identical and were distributed in blister packs. Scanning sessions occurred at the end of each regimen (varenicline 17.0 ± 4.2 days; placebo pill 16.5 ± 3.4 days). No washout interval separated medication periods. For those participants whose placebo regimen followed the varenicline regimen, carryover effects were assumed negligible given the ~24-hour elimination half-life of varenicline[5] and the fact that placebo varenicline scanning sessions and active varenicline scanning sessions were separated by more than 2 weeks.

For each of the pill conditions, nicotine or placebo patches were applied to the upper back at the start of fMRI visits (separated by 2.9 ± 1.7 days). Nonsmokers received a 7mg nicotine patch dose, while smokers received a dose that matched daily nicotine intake (21, 28, 35, or 42 mg patches for 10-15, 16-20, 21-25, and >25 cigarettes/day, respectively). The patch was worn for the duration of the 9-hour visit, which consisted of 2 MRI scans.

The PRL task was completed in the second of the 2 MRI scans and began approximately 6-7 hours after initial patch application. It was assumed that data collected within the 2-9 hour post-patch window was associated with steady plasma nicotine levels, in accordance with pharmacokinetic data.[6]

Before each scanning session, subjects were asked to abstain from alcohol for 24 hours, and to moderate caffeine intake for 12 hours. Smokers were required to abstain from cigarette use for 12 hours before and also during scanning days, but were not otherwise restricted on smoking behavior during the 6-8 week course of the entire study. At the start of each session, all participants underwent testing for recent drug and alcohol use, and expired carbon monoxide (CO) levels. CO levels of less than or equal to 15 parts per million (ppm) indicated abstinence. Medication dose was confirmed by the participant on the morning of the scanning session. Medication side effects and adherence were monitored by regular telephone assessments and at in-person visits.

#### Tasks

##### *Go-Nogo*

Participants were required to make a response on a button box when the stimulus presented on the screen (“X” or “Y”) was different from the previously presented stimulus (e.g. X-Y; Go trial) and suppress this prepotent response when the presented stimulus was identical to the previously presented stimulus (e.g. X-X; No-go trial). Stimuli alternated between “X” and “Y” on each trial, so the No-go stimulus was not always the same, ensuring that the task requires constant updating of the stimulus-response and cannot be solved with a simple visual search strategy [25]. Trials were spaced 1s apart, with the relatively infrequent Nogo trials temporally jittered to ensure optimal estimation of the BOLD response. Nogo trials were relatively rare (100 out of 1200 trials; 8.3%) to have roughly equal numbers of commission errors and successful inhibitions. Participants performed four 5-minute blocks of 300 trials each.

*Flanker*

Participants were asked to identify a target stimulus, an “H” or an “S” with a button press. At the start of each trial, prior to presentation of the target stimulus, flanker items (HH_HH or SS_SS) were presented. After 100ms, the center target stimulus was added to the array and both remained on the screen for 50ms. The design was event-related, allowing estimation of the BOLD response associated with each trial type. Trials were congruent (target stimulus matches flankers, 50% of trials) or incongruent (target stimulus does not match flankers, 50% of trials). A speeded version of the Flanker task with an individualized response deadline was used during the fMRI session to ensure an adequate difficulty level and error rate [36]. This individualized deadline was based on a practice run (130 trials) in a mock scanner, whereby the response deadline was taken to be the mean RT + 1SD on all correctly answered trials. When a participant missed this deadline, feedback indicated a missed response and encouraged faster responding on subsequent trials (no feedback was presented following either correct or error trials). Dependent measures were error rate, RT, and missed response rate per condition. Participants completed four 9-minute 130-trial runs with short rest periods between each.

*Flanker*

#### MRI acquisition

Whole-brain blood oxygenation level-dependent (BOLD) echo-planar images were acquired with a Siemens 3T Magnetom Allegra scanner (Erlangen, Germany). Thirty-three 5-mm-thick slices were acquired in the sagittal plane (TR = 2s, TE = 27ms, flip angle = 80°, field of view = 220 mm in a 64 × 64 matrix. For the Flanker task, 272 volumes were collected per run (four runs per session) and for the GNG task, 157 volumes were collected per run (four runs per session). Imaging data were collected with a delay (332ms) between volume collections to aid the processing of simultaneously recorded EEG data (not discussed here). Structural images were acquired using an MPRAGE sequence (TR = 2.5s; TE = 4.38ms; FA = 8°; voxel size = 1 mm3).

#### Small volume masks

Contrasts of interest were assessed within volumes of interest. The volumes of interest were defined by an OR mask of the control and error contrasts for each task across participants and sessions. Volumes were thresholded such that ~3,000 voxels were retained with minimum cluster sizes of 100 voxels. The maps were thresholded at $t > 10^8$ (mask size 3,436 voxels) and at $t > 10^3$ (mask size 2,708 voxels) for the Go NoGo task and the Flanker task respectively. The differences in threshold are due to design differences between the two tasks. The NoGo Contrast is estimated against the implicit baseline, which includes Go trials. This approach results in a much more robust map. Importantly, as the masks were defined on the basis of the average across all four sessions, they are unbiased in terms of the pharmacological manipulation. Circularity is therefore not a concern.

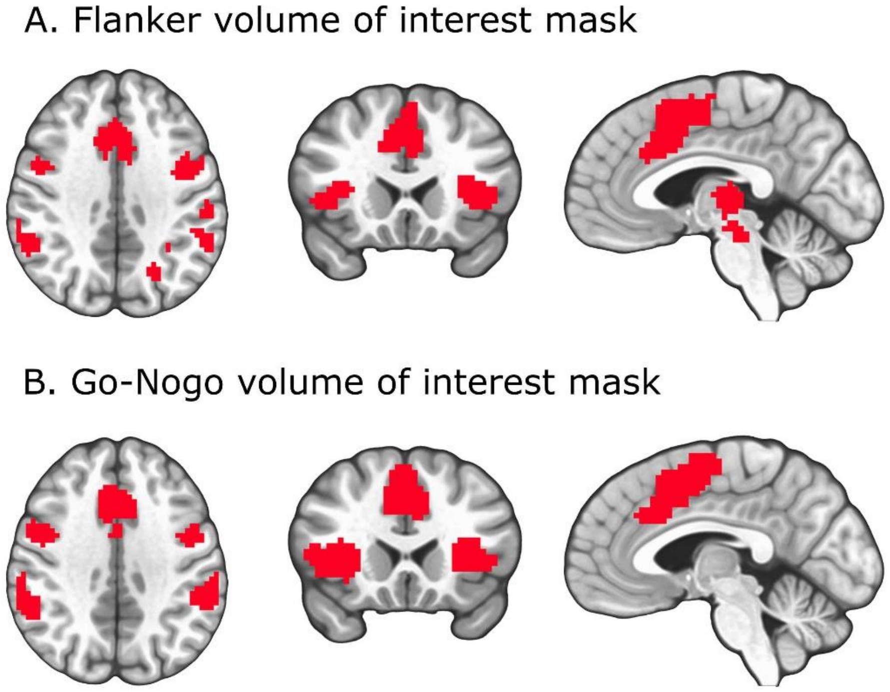


**Figure S2.** Small volume masks of interest for Flanker task (A) and Go-Nogo task (B). Radiological orientation: left is presented on the right.

### RESULTS

#### Demographics

Smoker and non-smoker groups were matched on gender and ethnicity. Smokers were moderately nicotine dependent (Fagerström scores: 5±2), smoked 18±8 cigarettes/day and reporting daily cigarette use for 18±11 years. Smokers were significantly older than controls ($t_{41.32}$=2.12, $p<0.05$), and age was therefore included as a covariate in behavioral and imaging between-group analyses. Data from one male non-smoker was excluded from all analyses due to consistently poor behavioral performance and excessive head motion.

Table S1. Demographics

| | smokers (N=24) | nonsmokers (N=20) | group differences |
|---|---|---|---|
| Gender (F/M) | 12 / 12 | 10 / 10 | t(42) = 0, p=1 |
| Age (mean +/- SD) | 35.8 +/- 9.9 | 30.4 +/- 7.2 | t(41.327) = 2.12, p=0.040 |
| IQ (mean +/- SD) | 105.95 +/-12.37 | 113.37 +/- 11.57 | t(37.0) = -1.93, p = 0.061 |
| race (AA/C/other) | 6 / 14 / 4 | 8 / 8 / 4 | F(3,40) = 0.49, p=0.6886 |
| Fagerstrom index | 5.00 +/- 1.9 | n/a | n/a |
| Years daily smoking | 18.0 +/- 10.6 | n/a | n/a |
| Cigarettes per day | 17.7 +/- 7.9 | n/a | n/a |

#### Behavioral results: Go NoGo

*Go-NoGo: behavioral results*

Table S2. Go-Nogo behavioral results

| | | NoGo Performance (prop) | | Go Performance (prop) | | Reaction Time (ms) | |
|---|---|---|---|---|---|---|---|
| | | Mean | SE | Mean | SE | Mean | SE |
| Non-smokers | Plac-Plac | 0.451 | 0.021 | 0.911 | 0.014 | 367.84 | 6.57 |
| | Plac-Nic | 0.476 | 0.019 | 0.951 | 0.007 | 366.75 | 4.45 |
| | Var-Plac | 0.471 | 0.019 | 0.958 | 0.010 | 366.20 | 3.79 |
| | Var-Nic | 0.448 | 0.021 | 0.938 | 0.009 | 366.29 | 5.02 |
| Smokers | Plac-Plac | 0.519 | 0.025 | 0.851 | 0.018 | 342.59 | 7.67 |
| | Plac-Nic | 0.536 | 0.021 | 0.950 | 0.010 | 346.19 | 5.56 |
| | Var-Plac | 0.526 | 0.023 | 0.897 | 0.010 | 355.95 | 7.22 |
| | Var-Nic | 0.536 | 0.018 | 0.941 | 0.012 | 355.73 | 5.03 |
| | Mean | 0.529 | 0.022 | 0.925 | 0.011 | 358.44 | 5.66 |

*Behavioral results: Flanker task*

Table S3. Accuracy and reaction time in the Flanker task

| | | Congruent correct (prop.) | | Incongruent correct (prop) | | Reaction Time (ms.) | |
|---|---|---|---|---|---|---|---|
| | | Mean | SE | Mean | SE | Mean | SE |
| Non-smokers | Plac-Plac | 0.856 | 0.016 | 0.643 | 0.035 | 442.18 | 6.21 |
| | Plac-Nic | 0.895 | 0.013 | 0.654 | 0.020 | 436.44 | 4.76 |
| | Var-Plac | 0.873 | 0.012 | 0.661 | 0.024 | 441.47 | 5.35 |
| | Var-Nic | 0.859 | 0.014 | 0.620 | 0.022 | 443.17 | 6.69 |
| Smokers | Plac-Plac | 0.789 | 0.017 | 0.599 | 0.022 | 437.94 | 3.71 |
| | Plac-Nic | 0.874 | 0.013 | 0.625 | 0.019 | 425.36 | 3.33 |
| | Var-Plac | 0.818 | 0.012 | 0.568 | 0.018 | 436.29 | 3.39 |
| | Var-Nic | 0.866 | 0.010 | 0.608 | 0.016 | 431.29 | 3.52 |
| | Mean | 0.837 | 0.013 | 0.622 | 0.022 | 436.77 | 4.62 |

Table S4. Missed responses in the Flanker task

| Missed Responses | | Congruent missed (prop.) | | Incongruent missed (prop.) | |
|---|---|---|---|---|---|
| | | Mean | SE | Mean | SE |
| Non-smokers | Plac-Plac | 0.081 | 0.014 | 0.126 | 0.021 |
| | Plac-Nic | 0.047 | 0.007 | 0.083 | 0.010 |
| | Var-Plac | 0.060 | 0.008 | 0.099 | 0.011 |
| | Var-Nic | 0.062 | 0.010 | 0.105 | 0.012 |
| Smokers | Plac-Plac | 0.132 | 0.016 | 0.191 | 0.020 |
| | Plac-Nic | 0.060 | 0.009 | 0.119 | 0.011 |
| | Var-Plac | 0.107 | 0.010 | 0.184 | 0.015 |
| | Var-Nic | 0.066 | 0.010 | 0.131 | 0.012 |
| | Mean | 0.077 | 0.011 | 0.130 | 0.014 |

#### Go-Nogo task map: corrections at p<0.001

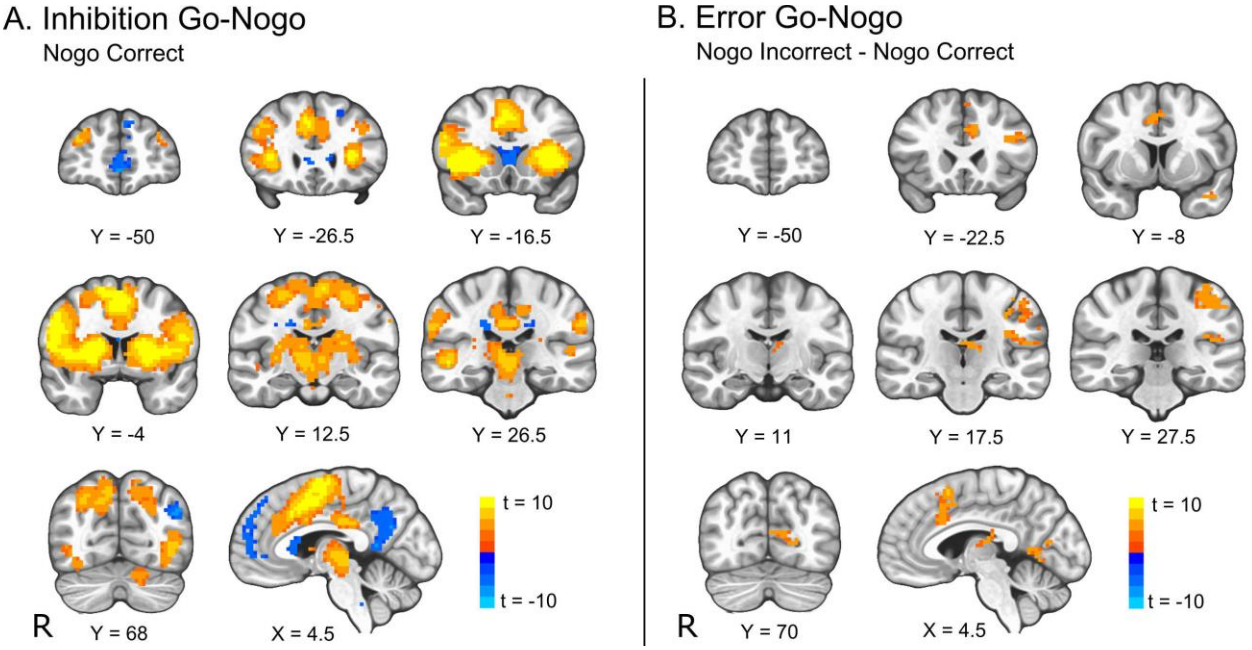


Figure S3. Task maps for the Go-Nogo contrasts. A. Brain areas with greater (orange) or smaller (blue) response to successfully inhibiting a prepotent response compared to baseline (correctly executed Go trials). B. Brain areas that showed greater (orange) response to failed inhibitions than to successful inhibitions on Nogo trials. Whole-brain corrected p<10-3 voxelwise, cluster size 22, FWE<0.05. Radiological orientation: right is presented on the left.

**Flanker task map: corrections at p<0.001**

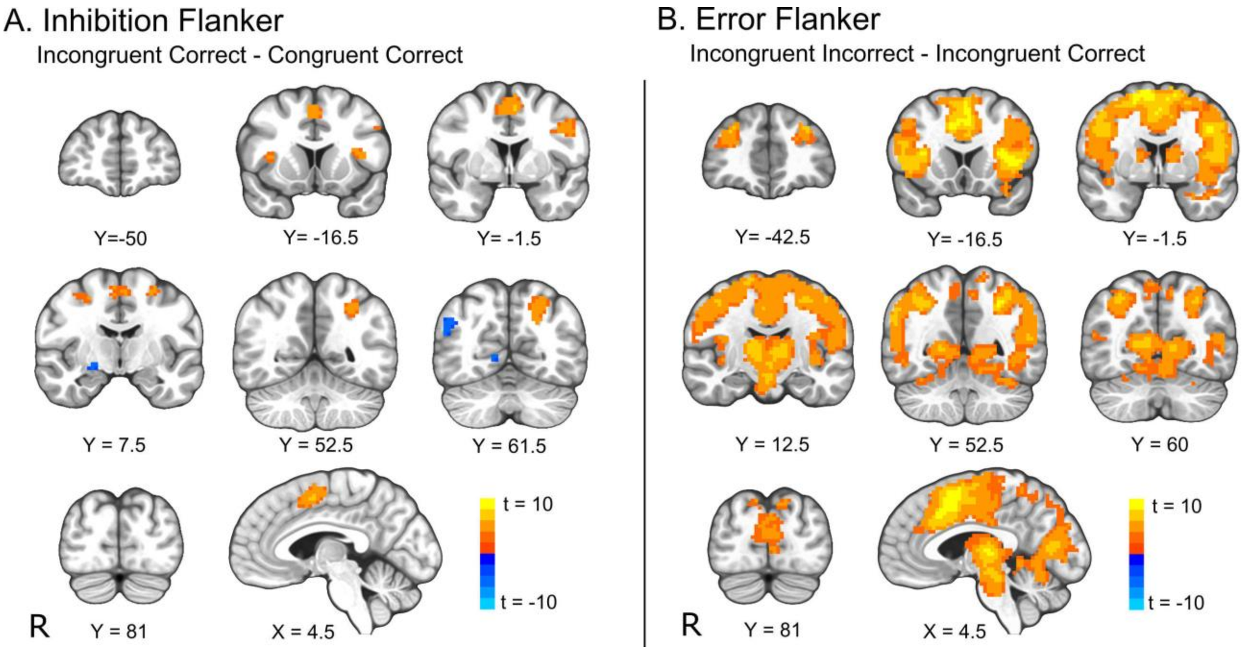


Figure S4: Task maps for Flanker task. A. Brain areas that showed more (orange) or less (blue) activity in response to correctly executed incongruent trials than to correctly executed congruent trials. B. Brain areas that showed greater activity during the incorrect execution of an incongruent trial than to the correct execution of an incongruent trial. Whole-brain corrected at FWE <0.05 (p<0.001 voxelwise, cluster size 22). Radiological orientation: right is presented on the left.
